# Reticulospinal drive increases maximal motoneuron output in humans

**DOI:** 10.1101/2022.01.24.477080

**Authors:** Jakob Škarabot, Jonathan P Folland, Aleš Holobar, Stuart N Baker, Alessandro Del Vecchio

## Abstract

Maximal rate of force development in adult humans is determined by the maximal motoneuron output, however the origin of the underlying synaptic inputs remains unclear. Here, we tested a hypothesis that the maximal motoneuron output will increase in response to a startling cue, a stimulus that purportedly activates the pontomedullary reticular formation neurons that make mono- and disynaptic connections to motoneurons via fast-conducting axons. Twenty-two men were required to produce isometric knee extensor forces “as fast and as hard” as possible from rest to 75% of maximal voluntary force, in response to visual (VC), visual-auditory (VAC), or visual-startling cue (VSC). Motoneuron activity was estimated via decomposition of high-density surface electromyogram recordings over the vastus lateralis and medialis muscles. Reaction time was significantly shorter in response to VSC compared to VAC and VC (i.e., the StartReact effect). The VSC further elicited faster neuromechanical responses including a greater number of discharges per motor unit per second and greater maximal rate of force development, with no differences between VAC and VC. We provide evidence, for the first time, that the synaptic input to motoneurons increases in response to a startling cue, suggesting a contribution of subcortical pathways to maximal motoneuron output in humans, likely originating from the pontomedullary reticular formation.

## INTRODUCTION

The rate of muscle force output is predominantly dictated by the central nervous system (Del Vecchio et al., 2019b; Dideriksen et al., 2020). Unlike in neonates that may modulate force output with a wide distribution of common synaptic input (Del Vecchio et al., 2020b), the maximal rate of force development in adult humans is determined by the speed of recruitment and discharge rate of motoneurons (Desmedt and Godaux, 1977; Van Cutsem et al., 1998; Del Vecchio et al., 2019b; Dideriksen et al., 2020). Compared to slower, sustained contractions which are typically characterised by the onion-skin phenomenon (De Luca and Erim, 1994), the discharge rate during rapid force production is substantially higher and peaks around the onset of force generation (Desmedt and Godaux, 1977; Del Vecchio et al., 2019b) followed by a non-linear decrease similar to in vitro spike frequency adaptation observed in rat (Sawczuk et al., 1995). Rapid contractions therefore allow insight into the maximal in vivo motoneuron recruitment speed and discharge rate, however the origin of inputs underlying maximal motoneuron output remains unclear.

Rapid generation of force production is a feedforward task with negligible influence of afferent feedback as evidenced by the recruitment of motoneurons before force onset (Del Vecchio et al., 2019b). Maximal motoneuron recruitment speed and discharge rate thus reflect the strength of the excitatory synaptic input (Duchateau and Baudry, 2014; Del Vecchio et al., 2019b). Supraspinal centres involved in motor command generation with connections to lower motoneurons seem likely substrates of characteristically strong synaptic input during rapid muscle force generation (Figure 1C). In addition to cerebral centres (e.g., pyramidal tract neurons descending from the primary motor cortex), subcortical neuronal populations such as the pontomedullary reticular formation play an important part in motor command generation (Buford and Davidson, 2004), the production of gross movement (Baker, 2011), and have been implicated in tuning locomotor speed (Capelli et al., 2017). Neurons in the reticular formation are large and fast-conducting (Brownstone and Chopek, 2018), with extensive branching of axons that make mono- and disynaptic connections with motoneurons (Riddle et al., 2009). Reticulospinal neurons are characteristically command neurons (Brownstone and Chopek, 2018), and have been shown to be involved in escape movement of vertebrate species where the need for rapid force production is required (Faber et al., 1989; Eaton et al., 2001).

**Figure 1.**
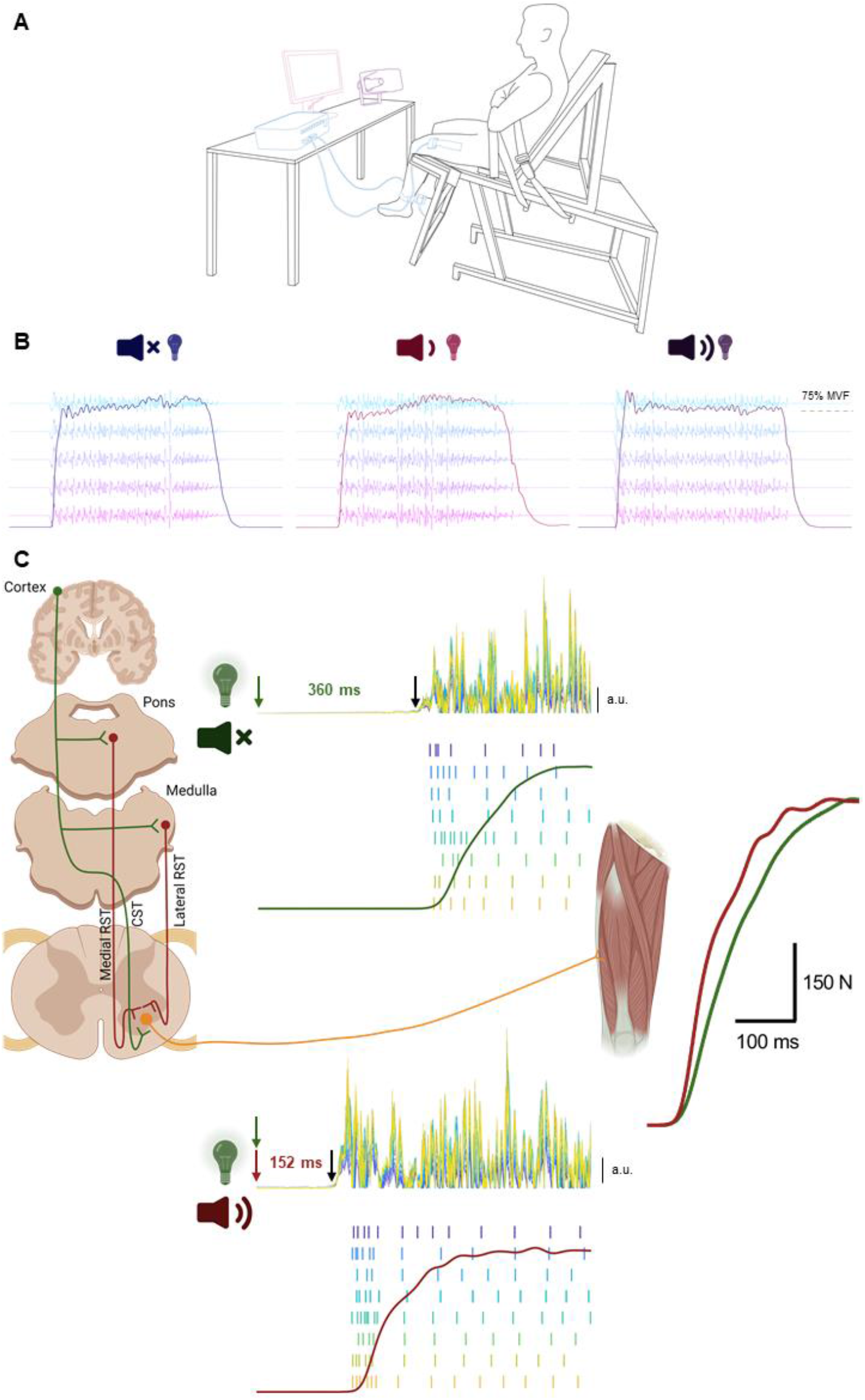
Experimental setup and descending pathways involved in the generation of the motoneuron output underpinning maximal rate of force development. A: Participants were seated in a custom-made isometric dynamometer whilst high-density EMG recordings were performed on vastus lateralis and medialis. Participants received either a visual (illumination of a light emitting diode, LED), visual-auditory (LED + quiet acoustic stimulus, 80 dB), or visual-startling (LED + loud acoustic stimulus, 110 dB) cue to which they had to respond by contracting the knee extensors “as fast and as hard” as possible. B: Examples of force and EMG recordings (recordings from only a single row of the high-density EMG grid placed on vastus lateralis are shown) in response to visual (left), visual-auditory (centre), and visual-startling cue (right). C: Presentation of a startling auditory stimulus (110 dB; red) purportedly activates neurons in pontomedullary reticular formation that transmit the activation signal to alpha motoneurons via the reticulospinal tract (RST), drastically shortening the reaction time (the time delay between stimulus delivery and visually determined onset of rectified EMG activity), increasing the initial discharge rate of motoneurons, and resulting in greater rate of force development.

Startling stimuli result in shortening of the reaction time (Valls-Solé et al., 1999) and augmented muscle force output (Anzak et al., 2011; Fernandez-Del-Olmo et al., 2014). These enhancements are thought to represent an involuntary release of pre-planned movement stored in subcortical circuits (Brown et al., 1991; Valls-Solé et al., 1995, 1999; Carlsen et al., 2003, 2004), likely in the pontomedullary reticular formation (Davis and Gendelman, 1977; Davis et al., 1982; Koch et al., 1992; Carlsen et al., 2004). As such, responses to startling stimuli have been used to infer reticulospinal contribution to human movement. For example, performance of gross motor actions that have at least a moderate reliance on reticulospinal input is accompanied by greater shortening of reaction time in response to startling stimuli compared to fine motor actions that have a greater reliance on corticospinal inputs (Carlsen et al., 2004; Baker and Perez, 2017; Tazoe and Perez, 2017).

In this study, adult humans were required to produce high isometric forces as rapidly as possible in response to presentation of a visual, visual-auditory, or visual-startling cue. We tested the hypothesis that the maximal motoneuron output will increase in response to a startling cue, a stimulus that purportedly activates neurons in the pontomedullary reticular formation, which will lead to more rapid generation of muscular force. Using a validated approach to high-density electromyography (HDsEMG) decomposition during rapid contractions (Del Vecchio et al., 2018, 2019b), we show greater rate of force development, and for the first time, greater number of discharges per motor unit per second in response to a startling auditory stimulus, suggesting reticulospinal contribution to maximal motoneuron output in humans.

## MATERIALS AND METHODS

### Participants

Twenty-two healthy, recreationally active men (mean ± SD; age: 24 ± 2 years, stature: 1.79 ± 0.07 m, mass: 80.9 ± 9.5 kg) volunteered to participate in this study. Participants were free from musculoskeletal or neuromuscular injury affecting function of the major joints of the lower limb, were not taking any medication known to affect the nervous system and reported no hearing-related impairments. The study was approved by Loughborough University Ethical Committee (2021-1749-3524) and was conducted in accordance with the Declaration of Helsinki except for registration in a database. Written, informed consent was provided by participants prior to taking part in any experimental procedures.

### Experimental design

Participants visited the laboratory on two occasions where they performed voluntary isometric knee extension contractions with their dominant limb. During the first session participants were familiarised with the experimental procedures by practicing maximal voluntary and rapid contractions (up to 10 trials and/or until satisfactory performance was reached; for criteria indicating valid trials, see ‘Force signal’ section of ‘Data processing and analysis’), and the StartReact protocol (for details, see below). Two to seven days later, participants returned to the laboratory for the experimental session. Participants were instructed to avoid any strenuous activity involving lower limbs within the 48 hours prior to the experimental session and caffeine consumption on the day of testing.

The experimental session started with a warm-up consisting of seven submaximal isometric knee extensions (3 × 50%, 3 × 75%, and 1 × 90% of perceived maximal voluntary force). Following warm-up, participants were instructed to perform a maximal effort isometric knee extension to estimate maximal voluntary force (MVF). Two trials were performed (60 seconds rest between trials) with strong verbal encouragement and live visual feedback of force level provided. If the two trials differed by more than 5%, an additional trial was performed. The highest instantaneous value was taken as MVF. Following MVF determination, participants performed up to five rapid submaximal practice trials in order to get used to producing the force as quickly as possible. After that, participants performed the StartReact protocol, adapted from Fisher et al. (2013), involving 18 rapid contractions in a block randomised order, with three blocks of six contractions. Participants were instructed to perform rapid contractions in response to illumination of a light emitting diode (LED; 20 ms) placed ∼1.5 metres in front of their participant. Out of six rapid contractions in a block, LED was presented either alone (visual cue, VC; two trials), with a quiet acoustic stimulus (visual-auditory cue, VAC; 80 dB, 500 Hz, 20 ms; RH40V, Adastra, Lisbum, UK; two trials), or a startling acoustic stimulus (visual-startling cue, VSC; 110 dB, 500 Hz, 20 ms; two trials), in a randomised order. Rapid contractions were separated by a random delay of 30-35 seconds (using customised scripts in Spike2 software; v9, Cambridge Electronics Design Ltd., Cambridge, UK). To avoid decrements in attention, participants were not required to maintain the focus on LED throughout the entirety of a block of contractions; rather they received a “get ready” command from the investigators ∼10 seconds before LED was due to be illuminated. Before commencing the rapid contractions, participants were presented with five consecutive startling acoustic stimuli every few seconds without performing any contractions for familiarisation with the startling cue (Fisher et al., 2013). For each rapid contraction, participants were instructed to contract “as fast and as hard as possible” to the target force of 75% of MVF in response to the ‘go’ signal (illumination of LED) and maintain the force level at 75% MVF target for ∼3 seconds. The ‘hold’ portion of the task was performed to increase the contraction duration for the purposes of HDsEMG decomposition algorithm which requires identification of sufficient number of independent sources (i.e., motor unit action potentials) from the electromyogram (Holobar et al., 2014; Del Vecchio et al., 2019b). Participants were additionally instructed to maintain a steady resting baseline force avoiding any countermovement or pre-tension (≤ 0.5 N) before the ‘go’ signal. Visual feedback of force was provided along with magnified feedback of the baseline force level (to facilitate trials without countermovement or pre-tension), and the first derivate of the force signal (as an indicator of the rate of force development achieved) was provided as performance feedback. The experimental set up along with an example of raw force and HDsEMG recordings is shown in Figure 1A and 1B, respectively.

### Experimental procedures

#### Force recordings

Participants were seated in a rigid custom-made isometric dynamometer designed to measure knee extensor forces. The knee and hip were flexed at 115 and 126°, respectively (180° = full extension). The selected knee angle has been previously shown to maximise isometric knee extension force production (Lanza et al., 2019). Participants were additionally strapped to the dynamometer across the chest and pelvis to prevent extraneous movement. The dynamometer utilised in this experiment has been shown to minimise joint angle changes during maximal isometric efforts (≤4°, compared to >15° of commercial dynamometers; Maffiuletti *et al*., 2016). The dominant leg was strapped to a metal brace placed behind the shank at ∼15% of tibial length (lateral malleolus to the knee joint centre) above the ankle, and in series with a calibrated S-beam strain gauge (Force Logic, Swallowfield, UK) positioned posterior and perpendicular to the tibia. The analogue force signal was amplified (×370), sampled at 2048 Hz, and acquired via a 16-bit multichannel amplifier (Quattrocento; OT Bioelettronica, Torino, Italy). The analogue force signal was also simultaneously sampled with a separate analogue-to-digital converter (2000 Hz, Micro 1401-3 and Spike2 v10 software, CED Ltd., Cambridge, UK) in order to display live force feedback to the participant, and to synchronise the recordings for calculation of the reaction time.

#### High-density electromyography

Multichannel, high-density surface electromyogram was recorded from vastus lateralis (VL) and vastus medialis (VM) muscles. Following skin preparation involving shaving, light abrasion, and cleaning with ethanol, semi-disposable grids of 64 electrodes (13 rows x 5 columns, 1 mm electrode diameter, 8 mm inter-electrode distance; GR08MM1305, OT Bioelettronica, Torino, Italy) were placed over the muscle bellies of vastus medialis and lateralis, with the long axis of the bi-dimensional grid oriented in line with the orientation of the muscle fibres as described previously (Martinez-Valdes et al., 2018). The placement of electrode grids on the skin was facilitated by disposable bi-adhesive foam layers (SpecsMedica, Battipaglia, Italy), the cavities of which were filled with conductive paste (AC Cream, SpecsMedica). A dampened strap ground electrode was placed on the ankle of the non-dominant leg, with reference electrodes (Kendall Medi-Trace, Canada) placed over the patella. High-density surface electromyogram signals were bandpass filtered (10-500 Hz), sampled at 2048 Hz, and recorded in a monopolar configuration via a 16-bit multichannel amplifier (Quattrocento; OT Bioelettronica, Torino, Italy) using OT Biolab+ software (OT Bioelettronica, Torino, Italy).

### Data processing and analysis

#### Force signal

During the offline analysis, the force signal was initially converted to force (N), and the baseline of the signal was gravity corrected. The force signal was then filtered with a zero-lag low-pass Butterworth filter with a cut-off frequency of 400 Hz. The onset of force was determined visually by a trained investigator using a systematic method (Tillin et al., 2010, 2013). Following determination of force onset, the force signal was additionally filtered by a zero-lag third order Butterworth filter with a cut-off frequency of 20 Hz, to eliminate high-frequency noise but maintain the undistorted force output (Del Vecchio et al., 2018). Trials were considered valid if no countermovement or pre-tension was displayed, and the force level was sufficiently high (≥75% MVF). Out of all trials, the three trials that were both valid and had the highest force at 150 ms after onset per the type of cue (VC, VAC, VSC) were selected for full analysis. The selection of the three best trials ensured that we captured maximal performance in response to specific cues. For the selected trials, the force signal was analysed in the first 200 ms following force onset; maximal rate of force development was calculated by the first derivate of force in each overlapping time window ranging from 1 to 200 ms (RFD0-X_max_, N.s^-1^) as described previously (Del Vecchio et al., 2019b). Such calculation of the first derivative by varying time windows allows for the assessment of the shifts in the maximal slope of the force-time curve among trials. Additionally, force was measured at 50, 100 and 150 ms following force onset, and the rate of force development was calculated as the first derivate of force for fixed time intervals from force onset to 50 ms, 50-100 ms, and 100-150 ms (Folland et al., 2014; Del Vecchio et al., 2018), in the interest of comparison with the literature. All variables were averaged across the three trials per the type of cue.

#### High-density electromyography

During off-line analysis (Matlab R2021a; Mathworks Inc., MA, USA), monopolar high-density EMG signals were initially band pass filtered (20-500 Hz) with a fifth-order, zero-lag Butterworth filter. Channels exhibiting poor signal-to-noise ratio, presence of movement artefact, poor skin-electrode contacts, or any irregularities were removed (typically ≤ 5% of all channels) using a semi-automated custom-made tool in Matlab based on area under the power spectrum and amplitude.

##### Decomposition

HDsEMG signals were decomposed into motor unit pulse trains (Figure 2A) via the Convolution Kernel Compensation algorithm (Holobar and Zazula, 2007). The validity of this algorithm has been repeatedly shown on both synthetic and experimental signals (Holobar et al., 2010, 2014). To improve decomposition accuracy, the 9 selected contractions (3 for each type of cue) were concatenated before decomposition. Concatenation was performed in a random order, with the order of concatenation stored for viewing only after all the analyses had been completed. The randomisation of concatenation ensured the investigator performing editing of motor unit pulse trains (Del Vecchio et al., 2020a) was blinded to the experimental conditions, thereby minimising investigator bias. To further optimise decomposition results, the editing process was performed on three contractions at the time, with motor unit spatial filters acquired during this process applied to the remaining contractions in overlapping windows (Francic and Holobar, 2021). Only motor units exhibiting a reliable discharge pattern with a pulse-to-noise ratio ≥ 30 dB (accuracy > 90%; false alarm rate < 5%) were retained (Holobar et al., 2014). To further test the validity of the decomposition, the motor unit action potential shapes (obtained via spike-triggered averaging) during the initial (transient force rise phase; first 10 discharges) and last 20 discharges (plateau phase of a contraction) were two-dimensionally cross-correlated (Figure 2C-D; Del Vecchio *et al*., 2019; Del Vecchio & Farina, 2020). An example of accurate identification of discharge timings following decomposition and further processing from one participant is demonstrated in Figure 2A-D.

**Figure 2.**
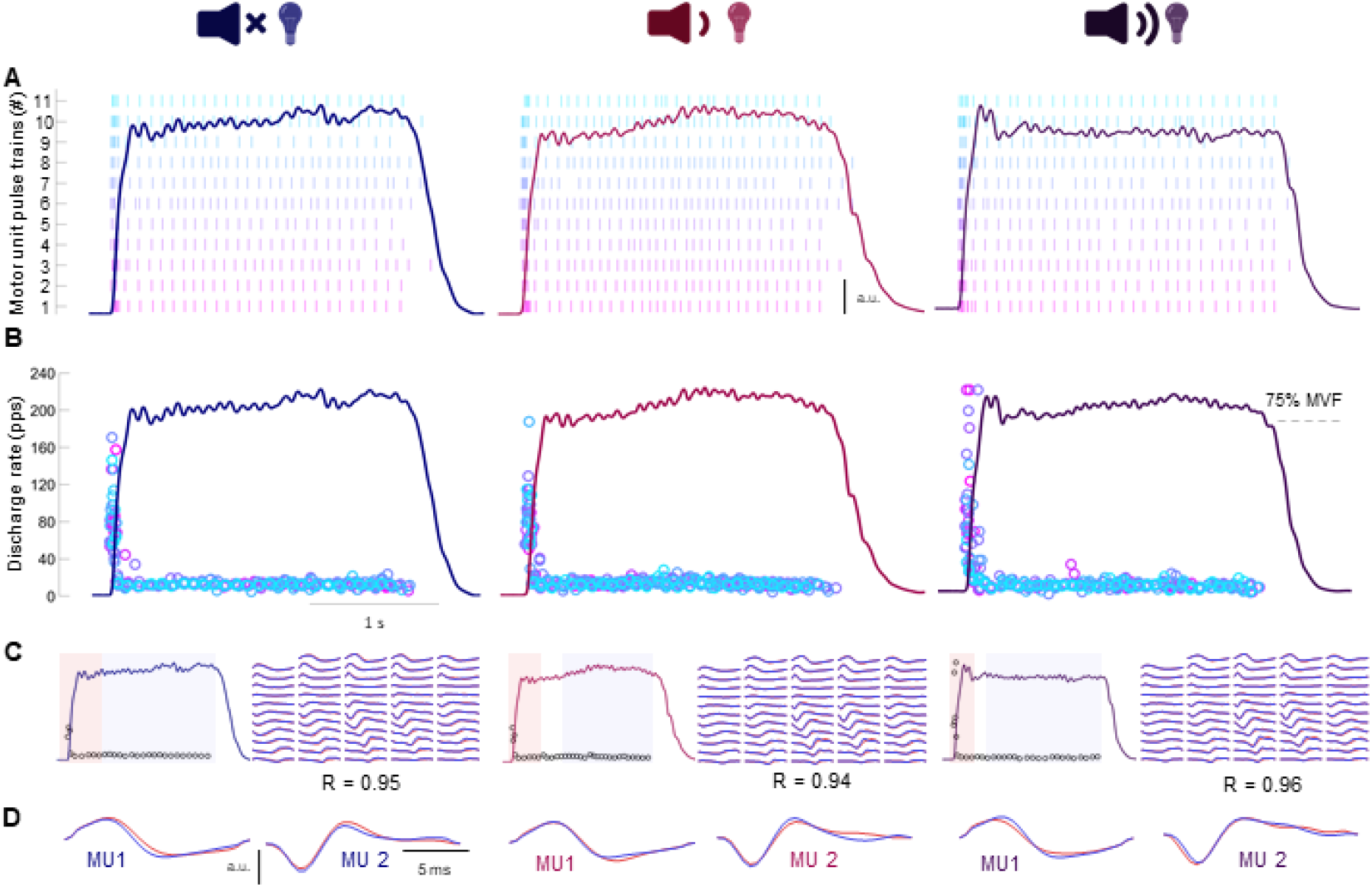
Decomposition of multichannel, high-density electromyogram. High-density surface electromyograms of vastus medialis obtained during rapid contractions in response to visual (LED only; left panel), visual-auditory (LED + 80 dB; centre panel), and visual-startling cue (LED + 110 dB; right panel) were decomposed into motor unit pulse trains (A), from which the discharge timings could be identified (B). To ensure validity of decomposition, spike triggered averaging was performed to compare the first 10 discharges (red; during transient force rise phase) to the final 20 discharges (blue; plateau phase) of individual motor units via two-dimensional cross-correlation (C). Note the high degree of similarity in motor unit action potential shapes of two example motor units both within- and between-contractions (D).

From the identified motor units several variables were computed. Maximal discharge rate of motor units was defined as the maximal instantaneous discharge rate at any point during the contraction, though this was very likely to occur within the initial few discharges. Discharge rate at recruitment was defined as the average discharge rate of the first five discharges, indicating neural drive around force onset, which has been shown to be highly associated with maximal rate of force development (Del Vecchio et al., 2019b). The strength of the neural drive was estimated by summing the discharge timings in a moving 35 ms epoch (corresponding to previously reported delay between onset of motor unit discharge and force), shifted every 1 ms up to 400 ms from the first discharge. The value of the summated discharge timings was then normalised to the time epoch (35 ms) and the number of active motor units, thus constituting the average number of discharges per motor unit per second (Del Vecchio *et al*., 2019). All variables were averaged across the three trials per type of cue.

##### Residual EMG activity

Whilst motoneuron recruitment has been shown to be one of the key determinants of rate of force development, we found only a single individual who exhibited discharges of an additional motor unit exclusively recruited in response to a startling stimulus (Figure 3). Considering that the current decomposition algorithms only allow discrimination of the activity of a limited proportion of the motor pool, with a bias towards neurons innervating superficial muscle fibres with large potentials (Farina et al., 2010), it is possible that neurons recruited exclusively in response to startling stimuli that contributed to augmented mechanical outcomes (see Results) were undetected.

**Figure 3.**
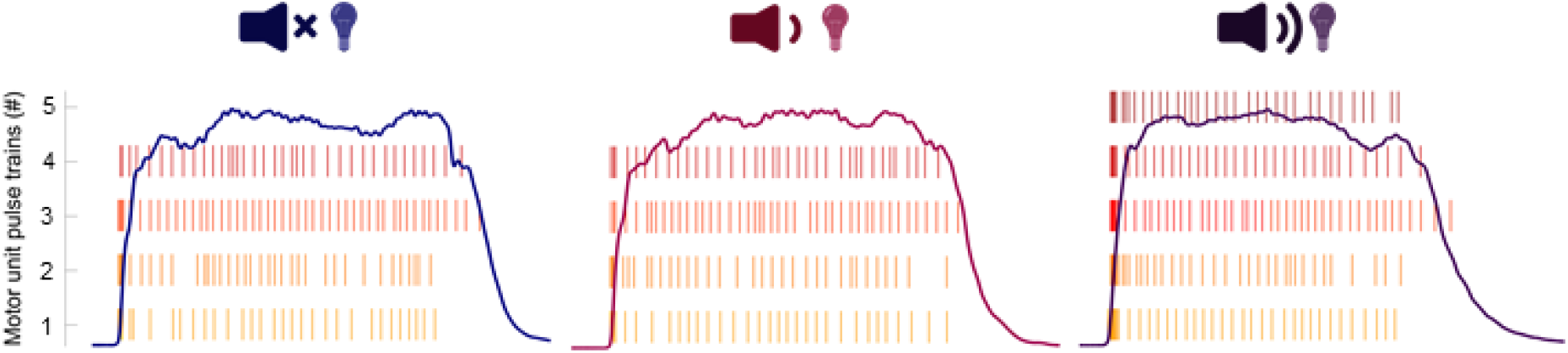
Example individual exhibiting additional motor unit recruitment in response to startling stimuli. An example participant in whom a visual-startling cue resulted in recruitment of an additional motoneuron (right panel) in vastus lateralis that was not recruited in response to visual (left panel) and visual-auditory cue (centre panel).

To check whether additionally recruited units might have been missed by decomposition, analyses of residual EMG activity were performed as follows. We first estimated motor unit action potentials by calculating spike triggered average of EMG signals, with the identified motor unit discharges used as triggers. After that, the motor unit action potentials of a given channel were convolved with the identified motor unit discharge patterns yielding the motor unit action potential train for each identified motor unit. For each EMG channel, the motor unit action potential trains of all the identified motor units were subtracted from the original EMG signals, yielding the residual EMG activity. This reflects the portion of the EMG signal which was not decomposed into identified unit. The root mean square of the residual EMG activity was then computed in a moving 35 ms epoch from EMG onset, shifted every 1 ms, and averaged across the three trials per type of cue. For the purpose of the statistical analyses, only the values of residual EMG activity in first 50 ms after force onset were compared across conditions as the root mean square EMG amplitude in this time window is most strongly associated with the rate of force development (Folland et al., 2014).

##### Reaction time

Because stimuli (visual, visual-auditory, or visual-startling) were delivered via a different system to the one with which EMG activity was recorded, the force signal that was simultaneously sampled and recorded in both (via two amplifiers) was initially resampled to the common sampling frequency of 2048 Hz, then afterwards cross-correlated to temporally align the recordings. From there, the reaction time was calculated as the delay between stimulus delivery and visually determined onset of rectified EMG activity of all recorded HDsEMG channels. The onset of rectified EMG activity was determined with the same systematic procedure used for determination of force onset. To confirm that the visual-startling stimuli evoked a specific response, the so-called reticulospinal gain was calculated as the difference in reaction times in response to visual and visual-startling cue, relative to the difference in reaction times in response to visual-auditory and visual-startling cue (Baker and Perez, 2017).

### Statistical analysis

Statistical analyses were performed using SPSS (v27; IBM, IL, US). Normality of data was assessed with the Shapiro-Wilk test. Reticulospinal gain and residual EMG activity were not normally distributed. A one-sample Wilcoxon signed rank test was therefore performed to assess whether the calculation of the reticulospinal gain was greater than 1. Residual EMG data was transformed with a common logarithm function to meet the assumption of normality. A one-way repeated measures ANOVA was performed to assess the differences in reaction times, maximal rate of force development, estimates of neural drive, discharge rate, and residual EMG activity between the three types of cues. A two-way (type of cue: VC, VAC, VSC; time: 0-50, 50-100, 100-150 ms) repeated measures ANOVA was performed to assess the differences in force and rate of force development in pre-selected time points/windows. A Greenhouse-Geisser correction was employed if the assumption of sphericity (Mauchly’s test) was violated. If significant F-values for main effects or interactions were found, analyses were continued with comparisons using least significant difference testing. Significance was set at an alpha level of 0.05. All data are presented as means ± standard deviation unless stated otherwise.

## RESULTS

### Reaction time

Reaction time was influenced by the type of cue in both VL (F_1.3,26.5_ = 87.5, p < 0.001; Figure 4A) and VM (F_1.4,30.0_ = 155.2, p < 0.001; Figure 4C), such that the fastest response was after the visual-startling cue compared to both visual-auditory (p < 0.001 for both VL and VM) and visual cues (p < 0.001 for both VL and VM). Furthermore, the reaction time in response to a visual-auditory cue was shorter than a visual cue (p < 0.001 for both VL and VM). Reticulospinal gain was significantly greater than 1 in both VL (Z = 4.1, p < 0.001; range 1.04 – 4.42; Figure 4B) and VM (Z = 4.1, p < 0.001; range 1.00 – 4.50; Figure 4D).

**Figure 4.**
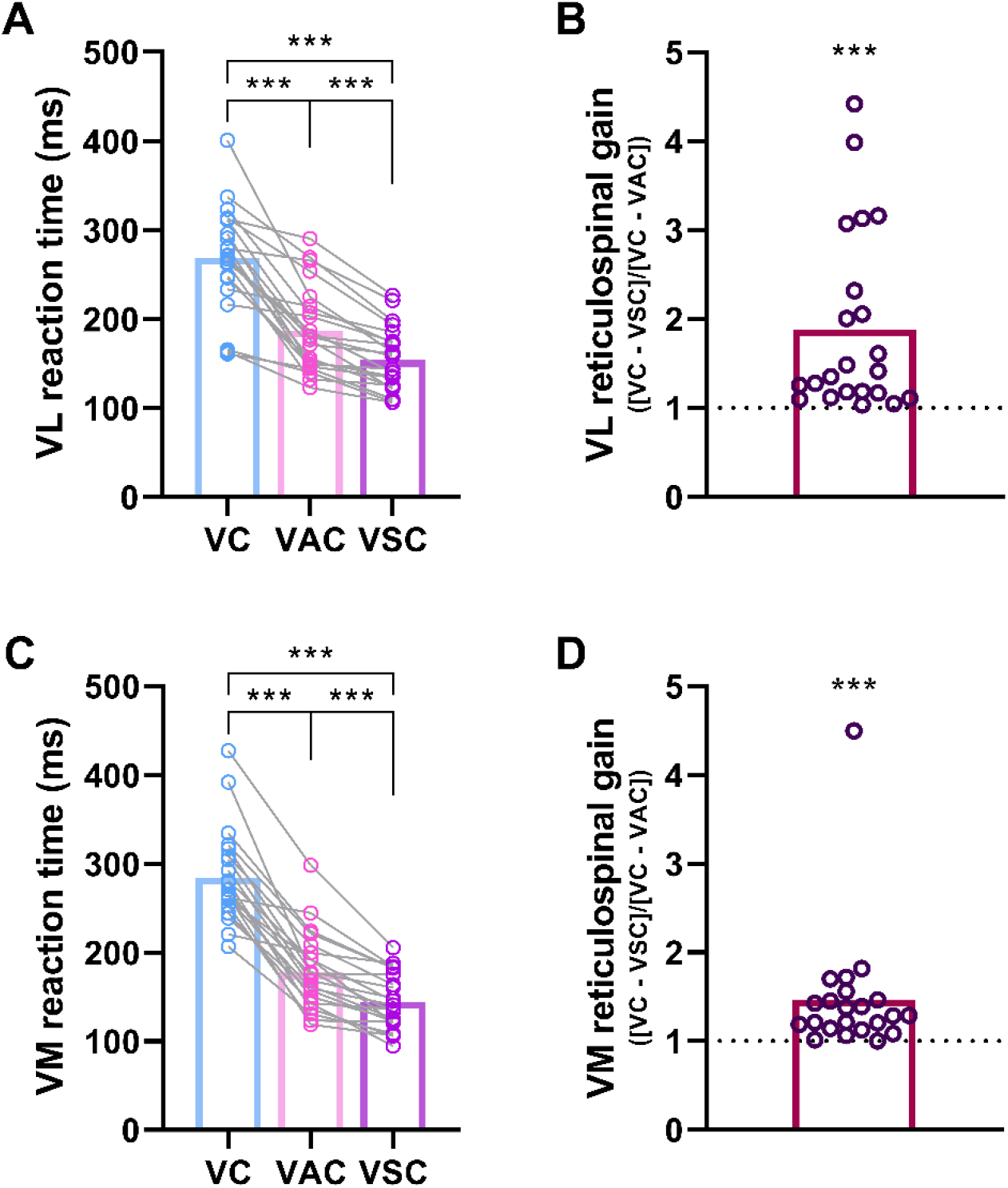
Reaction time and reticulospinal gain. Left panel: reaction times calculated as the time difference between the delivery of visual (VC; LED only), visual-auditory (VAC; LED + 80 dB), and visual-startling (VSC; LED + 110 dB) cues and the onset of EMG activity in vastus lateralis (VL; A) and vastus medialis (VM; C). N = 22; ***p<0.001 compared to other types of cues. Right panel: reticulospinal gain calculated as the difference in reaction time in response to visual and visual-startling cue relative to the difference in reaction time in response to visual and visual-auditory cue for the VL (B) and VM (D). N = 22; ***p<0.001 compared to 1.

### Mechanical output

There was an effect of the type of cue for absolute force at fixed time points following force onset (F_2,42_ = 29.5, p < 0.001), rate of force development during fixed time windows following force onset (F_2,42_ = 28.4, p < 0.001), and maximal rate of force development achieved during the entire force-time curve (F_2,42_ = 48.4, p < 0.001). Specifically, force production was on average ∼41, 18 and 14% greater at 50, 100 and 150 ms following force onset, respectively, when responding to the auditory-startling cue compared to visual and visual-auditory cue (p < 0.001 for all; Figure 5A), with no difference between the latter two type of cues at any time point (p ≥ 0.530). Similarly, rate of force development was greater in response to the auditory-startling cue compared to the other two types of cues in the 0-50 ms (∼33-49%; both p < 0.001) and 50-100 ms (∼9-13%; p ≤ 0.006) time window following force onset (Figure 5B), with no differences detected in the rate of force development between the visual and visual-auditory time cues during the same time windows (p ≥ 0.153). There were no differences in rate of force development when responding to different types of cues in the 100-150 ms time window (all p ≥ 0.279). Maximal rate of force development was ∼19% greater in response to the auditory-startling cue compared to visual and visual-auditory cues (both p < 0.001; Figure 5C), with no differences in maximal rate of force development between the latter cue types (p ≥ 0.358).

**Figure 5.**
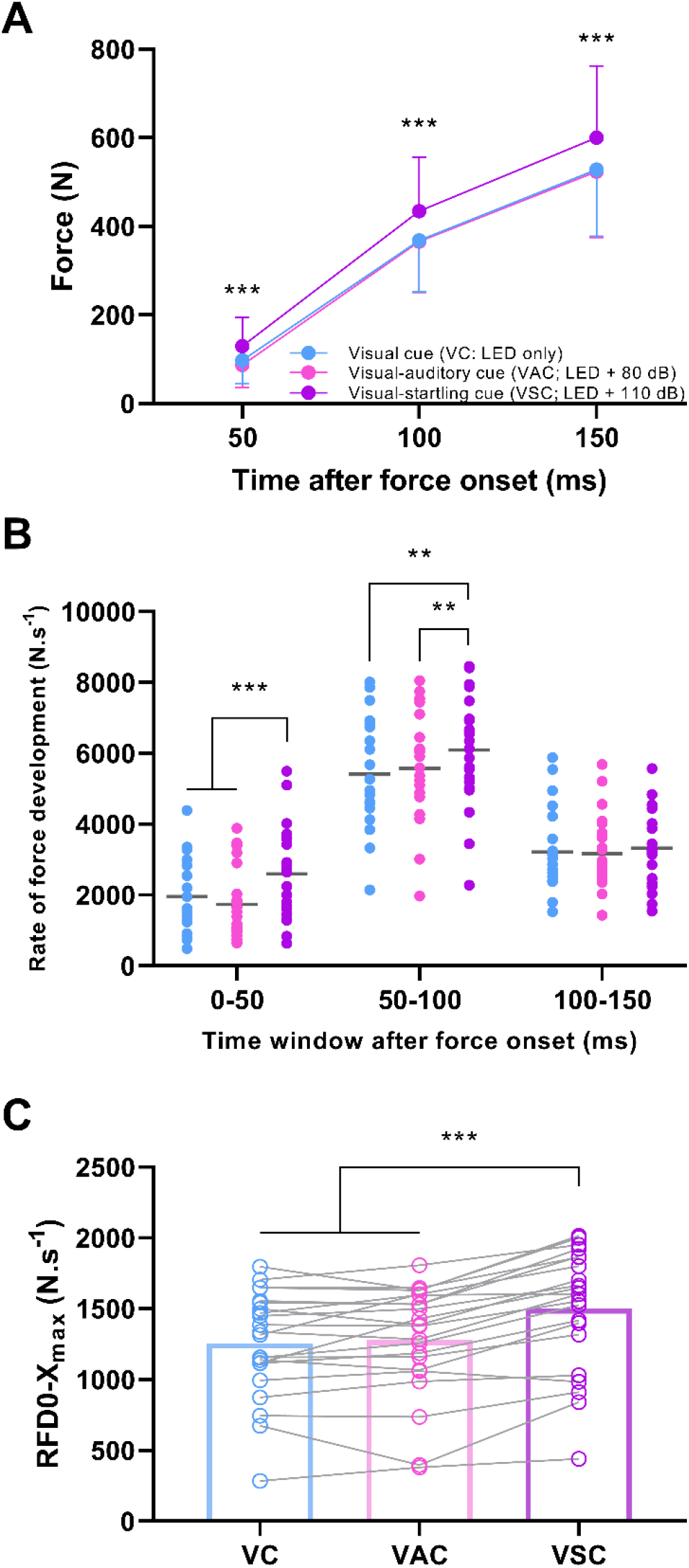
Force and rate of force development. Force produced at fixed time points following force onset (A), rate of force development (RFD; B) during fixed time windows following force onset (individual responses with means), and maximal rate of force development (C) achieved during the entire force-time curve (individual responses with means in bars) in response to either a visual cue (VC; LED only), visual-auditory cue (VAC; LED + 80 dB), or visual-startling cue (VSC; LED + 110 dB). N = 22; ***p<0.001, **p<0.010, *p<0.05 for visual-startling relative to other types of cues.

### Neural drive

Decomposition yielded reliable motor unit identification in 17 and 14 participants (out of a total sample of 22) in VL and VM, respectively. A total of 68 motor units were identified in the VL with the analysis yielding an average of 4.0 ± 2.2 motor units per individual (range 1-9). The average two-dimensional coefficient of correlation, indicating similarity of VL motor unit action potential shapes between the transient phase of force rise and the plateau region of the contraction, was 0.87 ± 0.07, with no differences detected during contractions in response to different types of cues (F_1.4,22.4_ = 2.1, p = 0.154). Within the VM, a total of 70 motor units were identified, with an average of 5.0 ± 3.8 motor units per respective contraction (range 1-11). The average two-dimensional correlation coefficient in the VM was 0.90 ± 0.08 and was not different during contractions in response to different types of cues (F_2,26_ = 1.2, p = 0.323).

The estimated neural drive, constituting the average number of discharges per motor unit per second, with respect to mechanical output is shown in Figure 6. The average value of the number of discharges per motor unit per second in the 50 ms from the first discharge was affected by the type of cue in VL (F_2,32_ = 16.4, p < 0.001; Figure 6A) and VM (F_1.3,17.2_ = 10.5, p = 0.002; Figure 6D), with the neural drive being greater in response to visual-startling cue compared to visual-auditory (VL: p = 0.002; VM: p = 0.020), and visual cue (VL: p < 0.001; VM: p = 0.007), consistent with the time window of the largest differences in mechanical output between cues (i.e. RFD within the first 50 ms after force onset). No differences in estimates of neural drive were detected between visual-auditory and visual cue (VL: p = 0.346; VM: p = 0.889).

**Figure 6.**
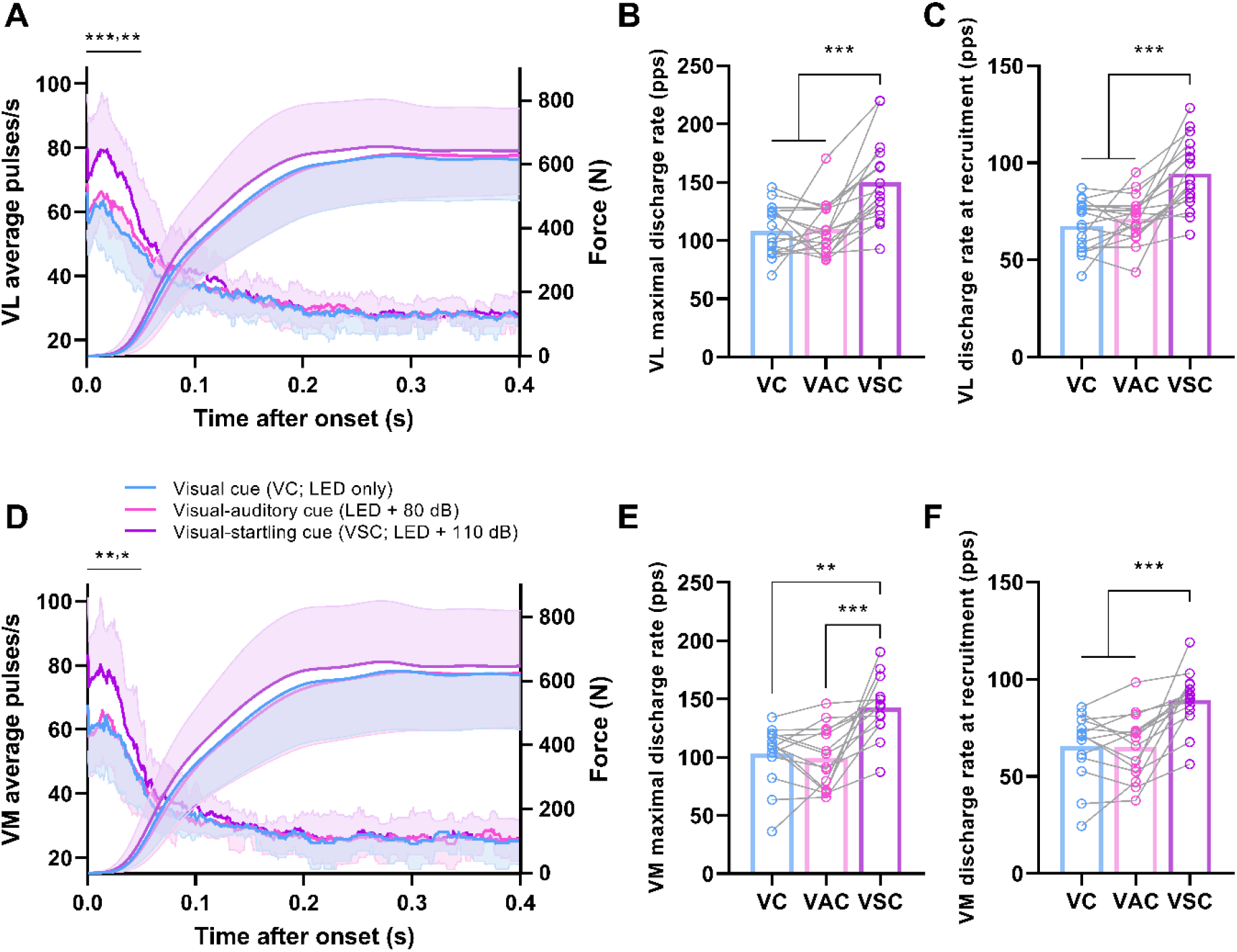
Motor unit discharge rate and force. The average number of discharges per motor unit per second, measured within a 35 ms moving window from the first discharge to 400 ms, in vastus lateralis (VL, N = 17; A) and vastus medialis (VM, N = 14; D), and the force output in response to a visual (LED only), visual-auditory (LED + 80 dB) and visual-startling (LED + 110 dB) cue from force onset. Shaded area represents standard deviation. Note the significant difference in the estimates of neural drive in response to the visual-startling cue relative to the other two types of cues in the first 50 ms, a time period corresponding to the greatest differences in rate of force development. Individual responses with means (bar plots) of the maximal instantaneous discharge rate (VL – B, VM – E) and the discharge rate at recruitment (mean of first five discharges; VL – C, VM - F) during rapid contractions in response to visual, visual-auditory, and visual-startling cue are also displayed. **p<0.001, **p<0.010, *p<0.05 for visual-startling relative to other types of cues.

The maximal discharge rate was affected by the type of cue in both the VL (F_2,32_ = 18.4, p < 0.001; Figure 6B) and VM (F_2,26_ = 14.9, p < 0.001; Figure 6E), and was greater in response to visual-startling compared to visual-auditory (p < 0.001 for both VL and VM) and visual cue (VL: p < 0.001; VM: p = 0.001), but no differences were detected between responses to the visual-auditory and visual cues (VL: p = 0.797; VM: p = 0.593). Similarly, discharge rate at recruitment, constituting the mean discharge rate during the first five discharges, differed in response to different cues in both VL (F_2,32_ = 23.7, p < 0.001; Figure 6C) and VM (F_2,26_ = 18.0, p < 0.001; Figure 6F). Specifically, the discharge rate at recruitment was greater in response to visual-startling compared to visual-auditory and visual cue (p < 0.001 for all). There were no differences detected in the discharge rate of the first five discharges between visual-auditory and visual cue (VL: p = 0.274; VM: p = 0.884). The discharge rate at the plateau phase of the contraction did not differ across conditions in both VL (17.3 ± 1.6 pps; F_2,32_ = 0.1, p = 0.917) and VM (17.5 ± 2.8 pps; F_2,26_ = 1.8, p = 0.196).

### Residual EMG activity

Residual EMG activity was affected by the type of cue in the VL in the first 50 ms from EMG onset (F_2,32_ = 8.6, p = 0.001; Figure 7A). Post hoc testing showed that the root mean square value of residual EMG signal was greater in response to the visual-startling compared to both the visual-auditory (p = 0.005) and visual cues (p < 0.001), with no differences between visual-auditory and visual cues (p = 0.902). However, there was no effect of cue on the residual EMG activity of the VM (F_1.5,23.3_ = 0.2, p = 0.756; Figure 7B). This analysis suggests that some additional units were likely recruited in response to the visual-startling cue, at least in VL, but these were not successfully decomposed by the algorithm.

**Figure 7.**
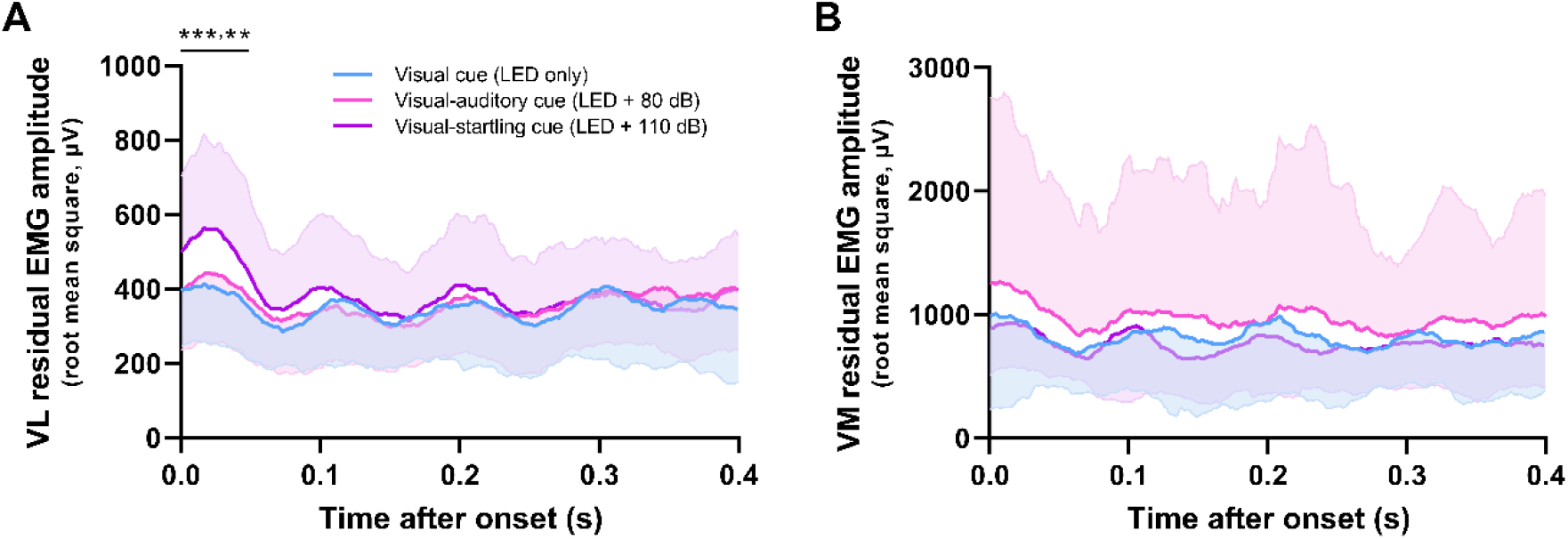
Residual EMG activity. Residual EMG was computed by subtracting motor unit action potential trains (obtained from the decomposition) from the original EMG signal. Root mean square amplitude of the residual EMG signal was then calculated with a moving 35 ms window from EMG onset for 400 ms in response to visual (LED only), visual-auditory (LED + 80 dB), and visual-startling cue (LED + 110 dB) for both VL (A) and VM (B). Statistical analyses were performed in the first 50 ms following onset due to the association of the EMG activity in this time window with the rate of force development. N = 17 for VL, N = 14 for VM. ***p<0.001, **p<0.010, *p<0.05 relative to other types of cues.

## DISCUSSION

In this study we tested a novel hypothesis that the maximal motoneuron output will increase in response to a startling cue, a stimulus that purportedly activates neurons in the pontomedullary reticular formation. We demonstrated that the presentation of a visual-startling cue shortened the reaction time and increased neural drive and the rate of force development. The increased synaptic input in response to a visual-startling cue suggests subcortical contributions to maximal in vivo motoneuron output in humans, likely originating from the pontomedullary reticular formation.

### Startling stimuli increase motoneuron output and rate of force development

A higher number of discharges per motor unit per second and greater rate of force development were only evident when responses to the visual-startling cue were compared to the visual and visual-auditory cues, with no differences between the latter two types of cues. Given that responses to a visual-auditory cue will be mediated by the cochlear nuclei and could thus result in intersensory facilitation (Nickerson, 1973), the observation that augmented neural drive and mechanical output were specific to the visual-startling stimulus further supports the notion that mediation occurred via increased descending input to motoneurons. Notably, the differences in neural and mechanical outputs were consistently the largest in the first 50 ms following respective onsets, which has been previously shown to be associated with descending neural mechanisms (Folland et al., 2014).

Startling stimuli have been shown to augment force output and increase the rate of force development (Anzak et al., 2011). However, the rate of force development and motoneuron output are related to the peak force achieved during the contraction (Desmedt and Godaux, 1977; Van Cutsem and Duchateau, 2005), thus the greater rate of force production reported previously (Anzak et al., 2011) might be an artefact of greater absolute force. In the present study, the contractions were performed from rest to the same high target force levels across all conditions, providing convincing evidence that the augmentation of the neural drive and the rate at which muscle force can be produced is a true function of a startling stimulus.

The discharge rate of motoneurons during rapid contractions was the highest at recruitment, with a large range of maximal instantaneous discharge rates between individuals as shown previously (Del Vecchio et al., 2019b). In the present study we extended these observations by demonstrating that the motoneuron discharge rate can be significantly enhanced when a startling stimulus is presented. These findings suggest that the neural substrate augmented by startling stimuli is directly responsible for high discharge rate that typically underpins force production during feedforward tasks. Given that startling stimuli purportedly activate neurons in the pontomedullary reticular formation (Davis et al., 1982; Koch et al., 1992; Carlsen et al., 2004), the increase in motoneuron discharge rate may reflect transformation of inputs transmitted via the reticulospinal pathway that makes both monosynaptic and disynaptic connections to motoneurons (Riddle et al., 2009).

Motoneuron recruitment has previously been shown to be a key factor in maximal rate of force development (Del Vecchio et al., 2019b; Dideriksen et al., 2020). However, we found only a single individual who exhibited discharges of an additional motor unit exclusively recruited in response to a startling stimulus (Figure 3), making it possible that higher-threshold neurons that contributed to augmented mechanical outputs in response to startling stimuli were largely undetected by decomposition. Such a hypothesis is consistent with the greater residual EMG amplitude in response to startling stimuli, at least in the vastus lateralis. Whilst this greater residual EMG amplitude may reflect a greater discharge rate of undetected units, global surface EMG amplitude has been shown to be more sensitive to changes in motor unit recruitment rather than discharge rate (Del Vecchio et al., 2017). Our analysis of residual EMG amplitude therefore suggests recruitment of higher-threshold motoneurons during the performance of a rapid, high-force contraction in response to a startling stimulus.

It has been speculated previously that rate of force development could be augmented as a result of increased synchronisation of motoneuron discharge rate (Semmler, 2002). Whilst it is possible that synchronisation of motoneurons was greater in response to a startling cue in the present study, such behaviour is more likely to be due to the intrinsic link between greater discharge rate and synchronisation, rather than a true augmentation of common synaptic input in adult humans (Del Vecchio et al., 2019a). Overall, our results are consistent with the notion presented previously that an increase in the rate of force development, facilitated by the startling stimulus in the present study, is a consequence of faster recruitment and greater discharge rate of motoneurons (Desmedt and Godaux, 1977; Van Cutsem et al., 1998; Del Vecchio et al., 2019b; Dideriksen et al., 2020), which seems to be, at least in part, driven by subcortical structures.

### The role of reticulospinal input in generating maximal in vivo motoneuron output

The response to a startling stimulus in humans differs from a startle reflex in that it does not habituate and does not exhibit pre-pulse inhibition (Valls-Solé et al., 2005). Randomisation of the order of cue presentation, along with highly consistent mechanical and neurophysiological outcomes in the present study make habituation and a learning effect an unlikely mechanism to explain our results. Several studies have suggested that the origin of an augmented response to a startling stimulus is subcortical, likely through the activation of neurons in the pontomedullary reticular formation (Davis et al., 1982; Koch et al., 1992; Carlsen et al., 2004), and we provide indirect support of this supposition via significant and specific shortening of the reaction time in response to a startling stimulus (i.e. significant reticulospinal gain; Baker and Perez, 2017). The reticulospinal neurons are characteristically command neurons (Brownstone and Chopek, 2018), with highly divergent postsynaptic connections that allow simultaneous innervation of many motor pools and thus facilitate synergistic execution of gross motor tasks (Peterson et al., 1979; Baker, 2011; Brownstone and Chopek, 2018; Zaaimi et al., 2018), which seems favourable for actions examined in the present study that required rapid force production of the quadriceps femoris. Indeed, previous studies have shown a greater StartReact response during gross motor tasks with a greater reliance on reticulospinal input compared to fine motor tasks (Carlsen et al., 2004; Baker and Perez, 2017; Tazoe and Perez, 2017).

Both healthy individuals after an intervention targeting the reticular formation (Germann and Baker, 2021), and patient populations with extensive cortical damage (Honeycutt and Perreault, 2012; Nonnekes et al., 2014; Choudhury et al., 2019) have shown an enhanced StartReact response. Nevertheless, the contribution of cortical influences to startling stimuli cannot be fully excluded (Marinovic and Tresilian, 2016), and similarly, the performance of a rapid, high-force task likely has a cortical component. For example, Baudry and Duchateau (2021) observed a specific response in the preparatory phase of rapid, compared to slower ramp contractions, including a steeper and delayed rise in corticospinal excitability, intracortical disinhibition, with only a limited increase in spinal excitability, suggesting a role of the primary motor cortex in encoding rapid contractions. It should be noted, however, that our results do not necessary conflict with those of Baudry and Duchateau (2021), but rather complement them. Indeed, there is an extensive network of cortico-reticular connections (Fregosi et al., 2017; Darling et al., 2018; Fisher et al., 2021; Figure 1C), thus in response to a startling stimulus, cortical inputs will likely be amplified by the reticulospinal neurons leading to a greater motoneuron output. Collectively, the present data, in conjunction with previous work, suggest that different neural substrates, both cortical and subcortical, likely act synergistically and contribute to fast recruitment and high discharge rate of motoneurons as well as their variability during the performance of rapid, high-force contractions.

## Conclusion

We demonstrated that when performing a rapid, high-force isometric task, presentation of a loud (startling) acoustic stimulus increased the average number of motor unit discharges per second and the rate of force production. Though cortical influences cannot be excluded, the increased synaptic input in response to a visual-startling cue suggests a subcortical contribution to maximal motoneuron output in humans, likely originating from the pontomedullary reticular formation.

## Acknowledgments

Dr Jakob Skarabot is supported by Versus Arthritis Foundation Fellowship (ref: 22569). Prof Ales Holobar is supported by Slovenian Research Agency (J2-1731, L7-9421, and P2-0041). Prof Stuart Baker is supported by the UK BBSRC (BB/V00896/X1). The authors thank Mr Jules Forsyth and Mr Apostolos Vazoukis for assistance with data collection and participant recruitment. The authors are also grateful to Mr Matic Škarabot for designing the schematic depicting experimental setup.

## Notes

**Conflict of interest:** The authors declare no competing financial interests.

### Competing Interest Statement

The authors have declared no competing interest.

